# A dual-retrieval-mode extinction procedure reduces fear renewal

**DOI:** 10.1101/2025.09.12.675473

**Authors:** Yingzhu Zeng, Zimeng Li, Hongbo Wang

## Abstract

A key limitation of exposure therapy is the return of fear after extinction. We drew on the memory-modulating benefits of selective retrieval and the learning-enhancing effects of variability to develop a novel dual-retrieval-mode extinction procedure. Forty-eight participants completed a four-day differential fear-conditioning paradigm comprising acquisition (60% reinforcement rate), two extinction sessions, and a renewal test. During the extinction phases, participants in the dual-retrieval group alternated between standard US-expectancy ratings (unrestricted retrieval) and safe/dangerous judgments in line with the outcome of the most recent CS+/CS− trial (restricted retrieval), whereas the control group completed standard expectancy ratings throughout extinction. Conditioned responding was assessed using skin conductance responses and subjective measures, including US expectancy, valence, and arousal ratings. Both physiological and subjective indices indicated that the dual-retrieval group exhibited reduced fear renewal relative to the control group. During extinction this group had shown larger skin conductance responses and higher US expectancy than the control group, but comparable arousal and valence ratings suggest this reflects reduced within-session habituation rather than stronger inhibitory learning. Because differential responding persisted, the benefit reflects enhanced competitive strength of the extinction memory rather than erasure of the fear memory. Manipulating retrieval demands during extinction may represent a promising strategy for reducing the fragility and context-dependence of extinction learning.

## 1. Introduction

### 1.1. Extinction Learning and the Challenge of Fear Relapse

Fear memories help humans avoid danger and are essential for survival. However, excessive and rigid fear can develop into anxiety and fear-related disorders such as post-traumatic stress disorder (PTSD), thereby compromising physical and mental health. Such maladaptive fear memories therefore need to be weakened or transformed into adaptive ones. Pavlovian fear conditioning is a classic laboratory paradigm for studying fear memory. In this paradigm, after a neutral stimulus (conditioned stimulus, CS) has been repeatedly paired with an aversive stimulus (unconditioned stimulus, US), individuals acquire a CS–US fear association, such that the CS presented alone elicits a fear response similar to that elicited by the US. This response constitutes conditioned fear. If, however, the CS is subsequently presented repeatedly without the US, the conditioned fear response gradually weakens; this phenomenon is known as fear extinction and constitutes the standard extinction procedure. Exposure therapy, which is grounded in the principles of fear extinction, remains one of the most effective treatments for anxiety and fear-related disorders (Carpenter et al., 2018; Craske, Treanor, Zbozinek, & Vervliet, 2022). However, extinction does not erase or modify the original CS–US association; rather, it establishes a new inhibitory CS–no US association. These two memories compete with each other, and the outcome of this competition determines whether fear is expressed (Bouton, 1993; Bouton & Moody, 2004; Myers & Davis, 2007). Substantial evidence indicates that fear returns under a variety of conditions, such as the passage of time (spontaneous recovery), a change of context (renewal), and exposure to a stressful event (reinstatement; Lonsdorf et al., 2017; Vervliet et al., 2013). Fear relapse is a major problem that exposure therapy must resolve, and it reflects a fundamental asymmetry: fear memories are robust and context-independent, whereas extinction memories are relatively fragile, context-dependent, and difficult to access under stress (Wicking et al., 2016). Enhancing the durability and accessibility of extinction memory is therefore an important goal for both basic and clinical research.

To date, researchers have developed a variety of strategies for enhancing extinction learning. Existing optimization strategies can be classified as fundamental (expectancy violation, attention toward the feared stimulus, removal of safety signals, mental rehearsal), advanced (deepened extinction, occasional reinforced extinction), and generalization-promoting (retrieval cues, multiple contexts, stimulus variability, positive affect; see Craske, Treanor, et al. (2022), for a review). Although these strategies rest on different theoretical frameworks, many of them share a common goal: modifying the way in which individuals process, retrieve, and update threat-related information during extinction, so as to promote more durable retention of extinction memory.

### 1.2. Memory Competition as the Core Obstacle in Extinction

At the outset of extinction, presentation of the CS first reactivates the consolidated CS–US fear memory. Because the expected US fails to occur, a prediction error arises, and this drives the formation of a new inhibitory CS–no US memory. Extinction is therefore a process in which retrieval and new learning occur simultaneously. As extinction proceeds, the CS accumulates two competing meanings: its original danger meaning (CS–US) and a newly acquired safety meaning (CS–no US). Both memory traces are activated by the CS and compete for retrieval. Because the fear memory is typically stronger and more consolidated, it proactively interferes with the encoding, consolidation, and retrieval of the extinction memory. This competitive interference renders the extinction memory fragile, context-dependent, and difficult to retrieve, increasing susceptibility to fear relapse after extinction. Based on this, Craske, Treanor, et al. (2022) proposed that state-of-the-art exposure therapy should simultaneously maximize inhibitory learning to consolidate the extinction memory and minimize retrieval of the original fear memory, and they identified these as the core mechanisms for achieving durable reductions in fear, anxiety, and avoidance. How, then, can extinction learning be maximized while retrieval of the fear memory is minimized?

### 1.3. Selective Retrieval as a Candidate Mechanism: Retrieval Practice and Retrieval-Induced Forgetting

Cognitive research has established that selective retrieval can alter the accessibility of competing memories, with two complementary consequences. First, items that have been retrieved become more recallable in the future than they would have been without practice. This benefit is referred to as the retrieval practice (RP) effect (Karpicke & Blunt, 2011; Roediger & Butler, 2011; Smith et al., 2016). Second, competing items that share a retrieval cue with the practiced items but are not themselves retrieved become less recallable, which is retrieval-induced forgetting (RIF; Anderson et al., 1994; Anderson & Hulbert, 2021; Murayama et al., 2014). The typical retrieval-practice paradigm comprises (1) a study phase, in which participants encode all of the items (e.g., Fruit–Orange, Fruit–Pear, Drink–Whisky, Drink–Brandy); (2) a retrieval-practice phase, in which participants selectively retrieve a subset of the studied items in response to a cue (e.g., Fruit–Or); these are the RP+ items, whereas unpracticed items from practiced categories (e.g., Fruit–Pear) are the RP− items and items from unpracticed categories (e.g., Drink–Whisky, Drink–Brandy) are the NRP items; and (3) a final test phase, in which participants recall all of the studied items. Recall of RP+ items typically exceeds that of NRP items, reflecting the RP effect. Critically, recall of RP− items falls below that of NRP items, reflecting RIF. This pattern has been interpreted as evidence that selective retrieval actively inhibits competing memory representations (Anderson, 2003). Given this dual function, selectively retrieving the safety-related extinction memory may simultaneously strengthen that memory trace (the RP effect) and suppress retrieval of the competing fear memory (the RIF effect), thereby providing an ideal mechanism for reducing fear relapse. Whereas previous research on the RP and RIF effects has been largely confined to the field of education, examining selective retrieval in the context of fear extinction extends the RP paradigm to a domain that differs both procedurally and ecologically, broadening its applicability to conditioned emotional responding.

### 1.4. Implementing Restricted Retrieval During Extinction: A 1-Back Design

A key methodological challenge is how to operationalize selective retrieval within the fear-conditioning paradigm. Cognitive studies have typically employed word-pair lists as learning materials (Murayama et al., 2014), whereas fear-conditioning studies use very simple pictorial or auditory stimuli that are associated with different event outcomes (threat or safety), yielding CS–US and CS–no US associations that lack an explicit verbal-categorical structure. To bridge these two paradigms, we adopted the guiding-sentence approach used by Glazier et al. (2021) in a social associative-learning task. That approach uses specific sentence prompts to direct participants toward a particular memory representation rather than allowing free retrieval across all encoded associations, successfully implementing restricted retrieval. Glazier et al. demonstrated that such directed retrieval selectively strengthened target memories while inducing forgetting of competitors that were not cued by the sentences, validating the approach for non-verbal associative content. Here, we extend this logic to the domain of conditioned fear by using sentence prompts to direct participants to selectively retrieve the safety outcome (absence of the US) of the most recent CS trial, analogous to the 1-back paradigm used in working memory research.

The specific design of these retrieval cue sentences was motivated by the principle of serial dependence in cognition. Perceptual judgments are strongly biased toward recent experience, a phenomenon that has been documented for orientation, numerosity, and emotion perception, and Bayesian modeling indicates that this bias improves processing efficiency by exploiting the natural temporal autocorrelation of environmental events (Cicchini et al., 2024; Ortega et al., 2023). This serial bias depends on conscious awareness of the preceding stimuli: Kim et al. (2020) demonstrated that only consciously perceived stimuli bias subsequent perceptual judgments, whereas suppressed stimuli do not. During standard extinction, the dominance of the fear memory and the negativity bias may prevent some individuals from consciously registering the absence of the US after the CS. Contingency-shift awareness, which here refers to the conscious registration of the transition from CS– US to CS–no US, is often lacking during extinction. Jovanovic et al. (2022) showed that this lack of awareness is associated with higher fear-potentiated startle responses and more severe PTSD symptoms. These findings suggest that the perceptual system’s natural tendency to weight recent experience can be harnessed by deliberately directing attention to the safety outcome of the most recent CS, thereby promoting the use of the most therapeutically meaningful CS–no US information.

In the present study, the guiding sentences instructed participants to predict the outcome of the current CS trial on the basis of the single most recent experience, which constitutes a 1-back retrieval pattern. Compared with standard extinction, this design offers several advantages. (1) It promotes retrieval of the CS–no US extinction memory while suppressing retrieval of the CS–US association. (2) It narrows the scope of memory search and facilitates extinction learning. During standard extinction, in order to predict the outcome of the current CS trial accurately, individuals may search the outcomes of several or even all preceding trials to compute the weights of the CS–US and CS–no US associations. (3) It increases certainty regarding the outcome. The outcome of the single most recent trial is entirely unambiguous, namely that no US occurred. However, in standard extinction, retrieving both the CS–US and the CS–no US associations renders the meaning of the CS ambiguous. (4) It promotes awareness of the shift in contingency from CS–US to CS–no US, facilitating adaptive updating when environmental contingencies change; as noted above, such awareness is associated with more effective fear reduction (Jovanovic et al., 2022).

### 1.5. Dual-Retrieval Mode Design: Integrating Restricted and Unrestricted Retrieval

During extinction, however, the 1-back retrieval pattern circumvents the prediction error (PE), which refers to the discrepancy between the expected and the actual outcome and is widely regarded as the primary driver of associative learning (Craske et al., 2014; Loock et al., 2025; Stemerding et al., 2023). To preserve the PE that drives extinction learning, the present study interspersed 1-back restricted retrieval trials within the unrestricted retrieval trials of standard extinction, creating a dual-retrieval mode. Under the unrestricted retrieval mode, participants provide US-expectancy ratings for the current CS trial, an index commonly used in human fear-conditioning research to monitor expectancy updating. Under the restricted 1-back retrieval mode, participants first recall the outcome (danger or safety) of the preceding CS+/CS− trial and then judge the outcome of the current CS+/CS− trial in a manner consistent with that recollection. This dual-retrieval design serves two functions. First, the standard extinction trials preserve the PE. Second, the alternation between retrieval modes introduces variability. Variability during learning is thought to enhance the memory strength of newly acquired information (Bjork & Bjork, 2006), and relative to massed or uniform practice it has been shown to produce superior long-term retention (Craske et al., 2014). The dual-retrieval mode therefore retains the memory-modulating benefits of selective retrieval while restoring the PE on which extinction learning depends and adding the learning-enhancing effects of variability.

### 1.6. The Present Study

We tested whether this dual-retrieval-mode procedure improves extinction outcomes. Specifically, we examined whether alternating between unrestricted and restricted 1-back retrieval during extinction enhances retention of the extinction memory and attenuates fear renewal, as indexed by skin conductance responses and by US-expectancy, valence, and arousal ratings. We predicted that participants trained under the dual-retrieval mode would show less fear renewal at test than participants trained under standard extinction. The procedure was evaluated as an integrated package: the aim was to determine whether the combination improves extinction outcomes, not to isolate the contribution of any single component (PE preservation, selective retrieval, or variability), and disentangling these mechanisms remains a task for future research. To our knowledge, this is the first study to implement selective retrieval directly within a fear-extinction paradigm. If the procedure proves effective, it would offer a novel and readily implementable means of optimizing exposure therapy.

## 2. Method

### 2.1. Participants

The required sample size was estimated using G*Power 3.1 (Faul et al., 2009), assuming a medium effect size of *f* = 0.25, a significance level of α = .05, and statistical power (1 − β) = 0.80, for either the two-way interaction of 2 (Group: Ctrl, DR) × 2 (Stimulus: CS+, CS−) or the three-way interaction of 2 (Group) × 2 (Stimulus) × 2 (Trial: Trial 1, Trial 2). This calculation indicated that at least 34 participants (17 per group) were required. Participants were recruited from the university community through posted advertisements and voluntary sign-up. A total of 52 right-handed undergraduate students (aged 18–21 years) were recruited. All had normal or corrected-to-normal vision and normal hearing. None had a history of physical illness or psychiatric disorder, and none had previously participated in a similar emotion experiment. Female participants were tested outside of their menstrual period to minimize hormonal influences on fear learning.

The study was conducted in accordance with the Declaration of Helsinki and was approved by the Ethics Committee of Henan University (approval number: HUSOM2019129; date of approval: 10 March 2019). Before the formal experiment, all participants signed a written informed consent form. Participants were informed of (1) the content of the assessments (including skin conductance recording, questionnaire completion, and subjective ratings); (2) the nature of the electrodermal recording and the fact that the electric shocks would be individually calibrated and strictly limited in voltage, such that they would not cause any harm; (3) the right to terminate and withdraw from the experiment at any time if they felt uncomfortable; (4) the confidentiality principle, namely that all data and information concerning them would be kept strictly confidential, and that they were likewise asked to maintain confidentiality regarding the content of the experiment; and (5) the monetary compensation provided to participants who completed all four days of the experiment, with a modest increase at the experimenter’s discretion according to the participant’s level of engagement.

Participants were subsequently randomly assigned either to the control (Ctrl) group, which received standard extinction (unrestricted retrieval mode), or to the dual-retrieval (DR) group, which received the dual-retrieval mode. Four participants withdrew during the experiment (three from the Ctrl group and one from the DR group). The final sample comprised 48 participants (7 male in total; 24 per group), a sample size consistent with that employed by Wang et al. (2021) in comparable fear-conditioning research. Sex was determined by participant self-report at the time of recruitment; no data on gender identity were collected.

### 2.2. Stimuli

Following previous research (Chen et al., 2021), the conditioned stimuli (CSs) were two pictures of cuboid objects that differed in color (red and yellow) (see **Fig**. **1A**). The two pictures, matched in luminance, were presented against either a black or a gray background, and appeared at the center of a 21-inch liquid crystal display. Each picture was presented for 4 s. For one of the stimuli, 60% of the trials were followed by the electric shock; this stimulus served as the CS+. The other stimulus was never followed by the shock and served as the CS−. The assignment of the two pictures to the CS+ and CS− roles was counterbalanced across participants.

The unconditioned stimulus (US) was a 200-ms electrical shock delivered to the inner surface of the left forearm, approximately 10 cm proximal to the wrist, via an STM200-1 stimulator module (BIOPAC Systems, Inc.) triggered by a parallel port output from E-Prime. Shock intensity was individually calibrated using an ascending staircase procedure: intensity was incrementally increased until the participant reported a discomfort rating of 8 on a scale ranging from 1 (comfortable) to 8 (extremely uncomfortable but bearable; Sperl et al., 2021). This individually determined intensity was then used throughout the experiment.

### 2.3. Measures

#### 2.3.1. Skin conductance response

Skin conductance was recorded continuously using a BIOPAC MP150 16-channel physiological amplifier (BIOPAC Systems, Inc., Goleta, CA, USA). A Skin Conductance Transducer (TSD203) connected to an EDA100C amplifier module was used for signal acquisition. TSD203 electrodes (6 mm contact diameter, 1.6 mm gel cavity depth; filled with GEL101A electrode gel, BIOPAC Systems, Inc.) were affixed to the distal palmar surfaces of the index and middle fingers of the participant’s left hand. Offline data processing was conducted using AcqKnowledge 4.2, the software supplied with the physiological recording system.

SCRs recorded during stimulus presentation were processed as follows. First, the raw skin conductance data were low-pass filtered at 1 Hz to remove noise. Second, responses to the CS+ and CS− in all phases and to the US during the fear acquisition phase on Day 1 were scored. For the US, the peak-to-trough difference (maximum minus minimum) in skin conductance within the 0.5–4.5 s window following US onset was taken as the raw unconditioned response (UR), and for the CS, the peak-to-trough difference within the 0.5–4.5 s window following CS onset was taken as the raw conditioned response (CR). Third, each participant’s raw CR values were divided by that participant’s mean raw UR to control for individual differences in electrodermal reactivity, and values smaller than 0.02 μS were scored as zero. Finally, the corrected data were square-root transformed to reduce distributional skew (Stussi et al., 2019; Wang et al., 2021).

#### 2.3.2. Subjective ratings

US expectancy ratings were collected as follows. To index explicit associative learning, participants rated their expectancy of receiving a shock following each CS presentation on a 9-point scale (anchored at 1 = “Certainly will not occur”, 5 = “Uncertain”, and 9 = “Certainly will occur”). The US expectancy prompt appeared immediately at CS offset (4 s after CS onset), and participants entered their response at their own pace by pressing the corresponding numerical key.

#### 2.3.3. Arousal and Valence

Participants also provided ratings of emotional valence (“Please rate your emotional feeling when viewing this image”; 1 = “Very pleasant” to 9 = “Very unpleasant”; O’Malley & Waters, 2018) and subjective arousal (“Please rate your emotional arousal when viewing this image”; 1 = “Very calm” to 9 = “Very fearful”) for each CS. Valence and arousal ratings were collected at the conclusion of the acquisition phase and both extinction phases, as well as on a trial-by-trial basis during the renewal test. All ratings were entered via the numerical keys (1–9) on the keyboard using the right hand.

### 2.4. Procedure

The experiment was conducted on four consecutive days and comprised a fear acquisition phase (Day 1), two extinction phases (Days 2 and 3), and a renewal test (Day 4). All procedures were programmed in E-Prime 2.0 (Psychology Software Tools, Pittsburgh, PA). Context was manipulated by changing the background screen color: Days 1 and 4 used a black background (RGB: 0, 0, 0), whereas Days 2 and 3 used a gray background (RGB: 128, 128, 128). The number of trials, the background, and the retrieval mode for each group in each phase are shown in **Fig. 1B**.

**Fig 1.**
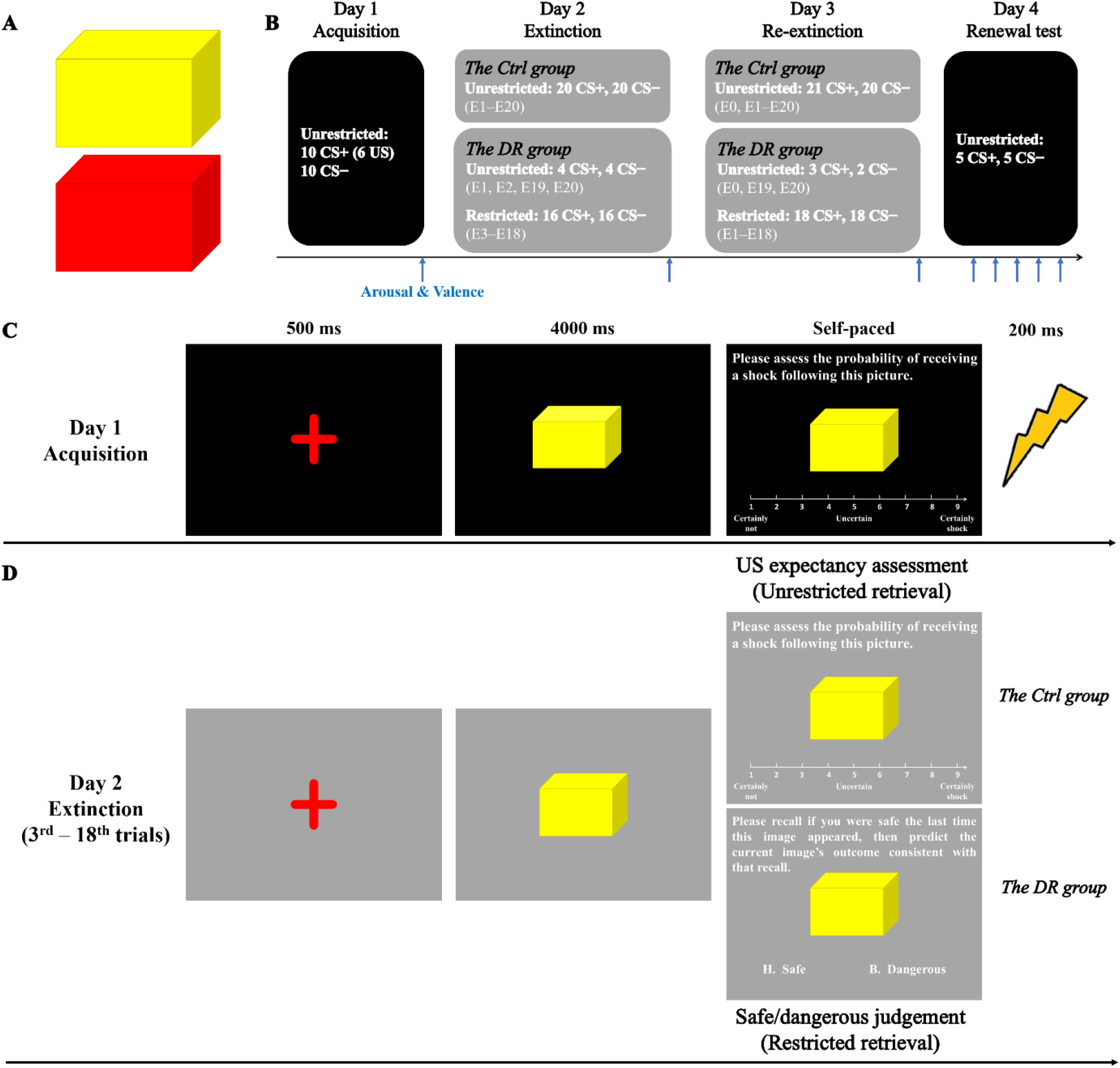
Experimental design and procedure (with the yellow cuboid depicted as the CS+ for illustrative purposes). (A) The conditioned stimuli (CS+ and CS−). (B) Schematic illustration of the task across four consecutive days. Blue arrows indicate the time points at which arousal and valence ratings were collected. (C) Schematic illustration of a trial during fear acquisition. (D) Schematic illustration of a trial during fear extinction. Two retrieval modes during extinction training: upper right, US expectancy rating (unrestricted retrieval); lower right, safe/dangerous judgment (restricted retrieval). E1 = the first trial of extinction, and so forth; E0 = the additional initial CS+ trial administered before re-extinction on Day 3.

#### 2.4.1. Day 1: Fear Acquisition

Upon arrival, participants signed the written informed consent form and provided demographic information before the experiment began. The experimenter attached the shock electrode to the inner surface of the participant’s left forearm, connected the physiological recording leads, and verified the settings of the shock stimulator. Shock intensity was individually adjusted for each participant (see section 2.2). Before the task began, participants had a 4–5 min rest period to allow SCR baseline levels to stabilize. Participants received the following instruction: “Two colored cuboid pictures will be presented. One of the pictures may be followed by an electric shock, whereas the other will never be followed by a shock. Please observe and discriminate between them carefully, and complete the ratings quickly as instructed.” The same instruction was provided on Days 2– 4, and the shock electrode was reattached at the beginning of each session. The CSs were presented in a pseudorandom order, with no more than two consecutive trials of the same CS+ or CS− within any experimental phase.

The acquisition phase comprised 10 CS+ trials (60% reinforcement rate) and 10 CS− trials presented in a pseudorandom sequence, with the first trial always being a reinforced CS+ trial. Each CS was presented for 4 s, after which the US expectancy rating prompt (“Please rate how likely it is that this picture will be followed by an electric shock”) appeared and participants responded by pressing a number key. On reinforced CS+ trials, the US was delivered immediately after the participant’s key press (Chen et al., 2024; see **Fig. 1C**). The intertrial interval (ITI) varied randomly between 11 and 16 s. At the end of the acquisition phase, participants completed valence and arousal ratings for the CS+ and the CS−.

#### 2.4.2. Day 2: Extinction

The extinction phase comprised 20 CS+ trials and 20 CS− trials presented in pseudorandom order, with the background color changed to gray. No US was delivered on any CS+ trial, and the ITI was the same as during acquisition. The two groups received different retrieval instructions on a subset of trials, which constituted the critical experimental manipulation (see **Fig. 1D**). The Ctrl group underwent standard extinction with unrestricted retrieval: US expectancy ratings were collected after every CS presentation across all 40 trials. We regard the US expectancy rating as an unrestricted retrieval mode because estimating an expectancy rating allows participants to integrate the outcomes of multiple preceding trials freely in order to compute the weights of the CS–US and CS–no US associations. In the DR group, the first two and the last two CS+ and CS− trials were identical to those of the Ctrl group and required US expectancy ratings, whereas on Trials 3 to 18 of each CS type the participants did not provide US expectancy ratings but instead completed a safe/dangerous judgment: “Please recall whether you were safe the last time this image appeared, and make a judgment about what will happen this time that is consistent with your recollection.” Participants indicated their judgment by pressing ‘H’ (Safe) or ‘B’ (Dangerous). This sentence-cued retrieval procedure prompted participants to selectively retrieve the CS–no US association from the most recent trial, an operation analogous to the 1-back paradigm used in working memory research. We refer to this 1-back retrieval as restricted retrieval because, under the sentence cue, the scope of memory search is restricted to the outcome of the immediately preceding trial. Keeping the initial and final trials identical to those of the Ctrl group enabled us to compare between-group differences in fear responding before and after the retrieval mode transition, while simultaneously preserving expectancy violation and increasing trial-to-trial variability. At the end of the extinction phase, all participants completed valence and arousal ratings for the CS+ and the CS−.

#### 2.4.3. Day 3: Re-extinction

The re-extinction procedure on Day 3 was largely identical to that of Day 2, with minor modifications. In order to increase the number of restricted retrieval trials, the two initial unrestricted retrieval trials of the DR group were also converted into restricted retrieval trials, so that only the final two CS+ and CS− trials remained identical to those of the Ctrl group and were used for between-group comparison. However, because the 1-back restricted retrieval procedure requires a preceding trial as a memory anchor, one additional initial CS+ unrestricted retrieval trial was administered at the beginning of re-extinction in the DR group to serve as the reference trial for restricted retrieval. To maintain procedural equivalence, one additional initial CS+ trial was likewise administered in the Ctrl group, with the structure of all remaining trials identical to that of Day 2. Both groups again completed valence and arousal ratings at the end of the phase.

#### 2.4.4. Day 4: Renewal Test

During the renewal test, the procedure was identical for the two groups. The background color of the CSs was restored to the black used during the acquisition phase. The test comprised 5 CS+ trials and 5 CS− trials. After each CS presentation, participants completed US expectancy, valence, and arousal ratings. At the end of the renewal test, all participants received monetary compensation and course credit.

### 2.5. Data Analysis

This study was not formally pre-registered. Apart from the manipulation checks reported in section 2.5.2, which were added after data collection and are labelled exploratory, the analytic approach was determined prior to data collection. All analysis decisions are reported in full below.

All data were analyzed using repeated-measures analyses of variance (ANOVAs) in JASP (version 0.95.4; JASP Team, 2025), with α = .05 as the criterion for statistical significance and effect sizes reported as partial eta-squared 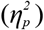. Where 95% confidence intervals are reported alongside an F test, they refer to the contrast in question (CS+ minus CS− for the main effect of Stimulus; Ctrl minus DR for effects involving Group), expressed in the units of the dependent variable rather than the effect size. When an interaction was significant, simple effects analyses were conducted, and Bonferroni-corrected p values were applied to control the error rate across multiple comparisons. Consistent with established conventions in the fear-conditioning literature (Dudziak et al., 2025), stronger fear recovery during the renewal test was operationally defined as (a) larger SCR amplitudes to the CS+, (b) higher US expectancy ratings for the CS+, and (c) higher arousal and more negative valence ratings, either specifically to the CS+ or, more generally, as indicated by a main effect of Group. The specific ANOVA models applied at each phase are described below.

#### 2.5.1. Acquisition and Extinction Phases (Days 1–3)

For the acquisition, extinction, and re-extinction phases (Days 1–3), SCRs and US expectancy ratings were computed as the mean of the final two trials for each stimulus type and submitted to a 2 (Group: Ctrl, DR) × 2 (Stimulus: CS+, CS−) repeated-measures ANOVA, with Group as a between-subjects factor and Stimulus as a within-subjects factor. Averaging across the final two trials provides a stable estimate of asymptotic responding within each phase while reducing the influence of single-trial variability. The CS arousal and valence ratings collected at the end of each phase were analyzed using the same Group × Stimulus model.

#### 2.5.2. Manipulation Checks

Compliance with the restricted retrieval instruction was assessed in the DR group. Because no US was delivered during either extinction phase, the correct judgment on every restricted retrieval trial was “safe”; accuracy was therefore defined as the proportion of “safe” responses. Accuracy was compared with the chance level of 50% using one-sample t tests, with Bonferroni correction applied across the four Day × Stimulus conditions, and was submitted to a 2 (Day: Day 2, Day 3) × 2 (Stimulus: CS+, CS−) repeated-measures ANOVA. To examine whether compliance changed across trials, the restricted retrieval trials of each phase were divided into early and late halves (8 trials each on Day 2, 9 trials each on Day 3) and entered as an additional within-subjects factor. Because accuracy approached the ceiling in many participants, the Day and Stimulus effects were re-examined using Wilcoxon signed-rank tests. Response times on the restricted retrieval trials were computed for correct trials only and are reported descriptively.

#### 2.5.3. Renewal Test (Day 4)

Fear renewal on Day 4 was evaluated using a 2 (Group: Ctrl, DR) × 2 (Stimulus: CS+, CS−) × 2 (Trial: Trial 1, Trial 2) repeated-measures ANOVA. Only the initial two trials were analyzed, on the basis of the following considerations. First, the impact of a context shift on conditioned fear responding is typically greatest at the beginning of the test and diminishes rapidly as the new context is learned; the initial trials therefore provide the most sensitive window for detecting fear renewal. Second, averaging across the first two trials reduces the influence of single-trial noise while remaining within the window in which the impact of the context shift is maximal, and retaining Trial as a factor allowed us to check whether any group difference was stable across the initial trials of the test. Third, given that after two consecutive days of extinction participants show rapid extinction or habituation to the non-reinforced CS+, including additional later trials would attenuate and confound the relapse signal.

In the present study, fear responding during the renewal test was not compared with responding at the end of the preceding extinction phase. This decision was based on several methodological considerations. Residual fear at the end of an extinction phase reflects the combined influence of extinction learning and within-session habituation, both of which are difficult to disentangle and may affect the two groups differently (Brown et al., 2017; Craske et al., 2014; Pace-Schott et al., 2014). Because the DR group alternated between unrestricted, restricted, and unrestricted retrieval, whereas the Ctrl group used unrestricted retrieval throughout, the DR group would be expected to show less habituation; group differences at the end of (re-)extinction might therefore reflect differences in habituation rather than in extinction learning. In addition, the first trial of each new phase typically elicits an orienting response in skin conductance that increases SCR amplitude independently of conditioned fear. Comparing the final trial of re-extinction with the initial trial of the renewal test would therefore introduce multiple confounds, including habituation, orienting responses, and context change, all of which would greatly reduce the interpretability of the results. The analyses were consequently restricted to comparisons within the test phase.

## 3. Results

Data from all experimental phases (acquisition, extinction, re-extinction, and the renewal test) are presented in **Fig. 2**.

### 3.1. Fear Acquisition

SCRs were averaged across the final two CS+ and CS− trials and submitted to a 2 (Group: Ctrl, DR) × 2 (Stimulus: CS+, CS−) repeated-measures ANOVA. The main effect of Stimulus was significant (CS+ > CS−), *F*(1, 46) = 24.32, *p* < .001, 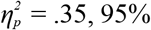CI [0.08, 0.19]. The main effect of Group was not significant, *F*(1, 46) = 0.73, *p* = .397, nor was the Group × Stimulus interaction, *F*(1, 46) = 0.73, *p* = .397.

The same Group × Stimulus repeated-measures ANOVA was applied to the mean US expectancy ratings across the final two CS+ and CS− trials, and to the arousal and valence ratings for the CS+ and CS− collected at the end of fear acquisition. The results showed that the CS+ elicited significantly stronger fear responses than the CS− on all subjective measures [US expectancy: *F*(1, 46) = 114.04, *p* < .001, 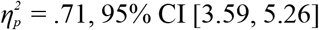; arousal: *F*(1, 46) = 85.76, *p* < .001, 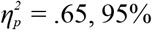CI [2.56, 3.98]; valence: *F*(1, 46) = 80.05, *p* < .001, 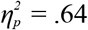, 95% CI [2.44, 3.85]]. The main effect of Group was not significant for any of the subjective measures [US expectancy: *F*(1, 46) = 1.50, *p* = .227; arousal: *F*(1, 46) = 0.70, *p* = .408; valence: *F*(1, 46) = 0.01, *p* = .935], nor were the Group × Stimulus interactions [US expectancy: *F*(1, 46) = 0.001, *p* = .983; arousal: *F*(1, 46) = 0.003, *p* = .953; valence: *F*(1, 46) = 0.17, *p* = .680].

Taken together, the SCR and subjective rating results indicate that the two groups acquired conditioned fear to a comparable degree.

**Fig 2.**
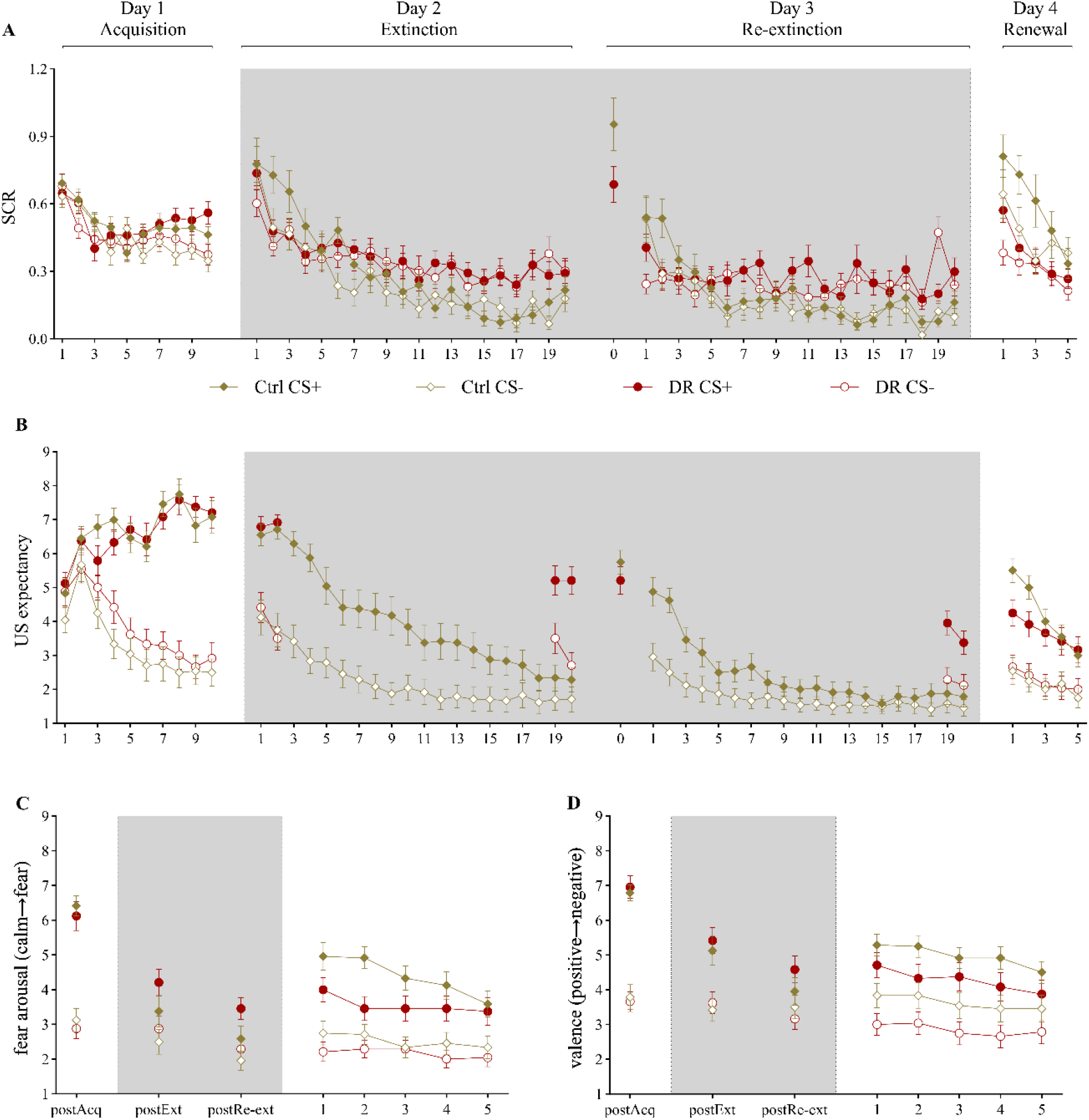
Dual-retrieval-mode extinction training reduces fear renewal. (A) SCRs for each trial across the four phases. (B) US expectancy ratings for each trial across the four phases. (C) Arousal ratings at the end of acquisition, extinction, and re-extinction, and for each trial of the renewal test. (D) Valence ratings at the end of acquisition, extinction, and re-extinction, and for each trial of the renewal test. SCR = skin conductance response; US = unconditioned stimulus; CS = conditioned stimulus; Ctrl = the control group; DR = the dual-retrieval group; “0” on Day 3 = the reference trial for restricted retrieval. Error bars represent standard errors.

### 3.2. Extinction

SCRs averaged across the final two CS+ and CS− trials were submitted to a Group × Stimulus repeated-measures ANOVA. The main effect of Group was significant, *F*(1, 46) = 7.84, *p* = .007, 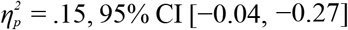, with higher SCRs in the DR group than in the Ctrl group. The main effect of Stimulus was not significant, *F*(1, 46) = 0.02, *p* = .877, nor was the Group × Stimulus interaction, *F*(1, 46) = 2.28, *p* = .138.

The same repeated-measures ANOVA was applied to the mean US expectancy ratings across the final two CS+ and CS− trials, and to the arousal and valence ratings for the CS+ and CS− collected at the end of extinction.

For US expectancy, the main effect of Group was significant, *F*(1, 46) = 27.25, *p* < .001, 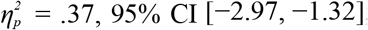; the main effect of Stimulus was significant, *F*(1, 46) = 18.20, *p* < .001, 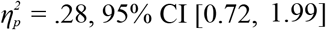; and the Group × Stimulus interaction was also significant, *F*(1, 46) = 5.58, *p* = .022, 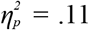. Simple effects analyses of the interaction indicated that differential responding (CS+ > CS−) remained significant in the DR group, *p* < .001, 95% CI [1.20, 3.01], whereas no significant difference between the CS+ and the CS− was observed in the Ctrl group, *p* = .185. In addition, US expectancy ratings were significantly higher in the DR group than in the Ctrl group for both the CS+, *p* < .001, 95% CI [−3.96, −1.83], and the CS−, *p* = .009, 95% CI [−2.42, −1.37]. For arousal, the main effect of Stimulus was significant (CS+ > CS−), *F*(1, 46) = 16.62, *p* < .001, 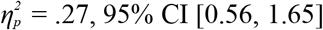, whereas the main effect of Group was not significant, *F*(1, 46) = 1.63, *p* = .208, and the Group × Stimulus interaction was not significant, *F*(1, 46) = 0.72, *p* = .402. For valence, the main effect of Stimulus was significant (CS+ > CS−), *F*(1, 46) = 25.15, *p* < .001, 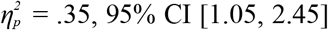, whereas the main effect of Group was not significant, *F*(1, 46) = 0.45, *p* = .506, and the Group × Stimulus interaction was not significant, *F*(1, 46) = 0.01, *p* = .905.

### 3.3. Re-extinction

SCRs averaged across the final two CS+ and CS− trials were submitted to a Group × Stimulus repeated-measures ANOVA. The main effect of Group was significant, *F*(1, 46) = 13.32, *p* < .001, 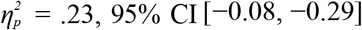, and the Group × Stimulus interaction was also significant, *F*(1, 46) = 4.46, *p* = .040, 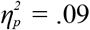, whereas the main effect of Stimulus was not significant, *F*(1, 46) = 3.01, *p* = .090. Simple effects analyses of the interaction indicated that SCRs were significantly higher in the DR group than in the Ctrl group for both the CS+, *p* = .027, 95% CI [−0.25, −0.02], and the CS−, *p* < .001, 95% CI [−0.37, −0.13]. Within-group comparisons showed no significant difference between the CS+ and the CS− in the Ctrl group, *p* = .791, whereas in the DR group SCRs to the CS− were higher than SCRs to the CS+, *p* = .009, 95% CI [−0.19, −0.03].

The same repeated-measures ANOVA was applied to the mean US expectancy ratings across the final two CS+ and CS− trials, and to the arousal and valence ratings for the CS+ and CS− collected at the end of re-extinction. For US expectancy, the main effect of Group was significant, *F*(1, 46) = 9.36, *p* = .004, 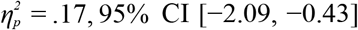; the main effect of Stimulus was significant, *F*(1, 46) = 30.44, *p* < .001, 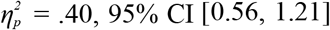; and the Group × Stimulus interaction was also significant, *F*(1, 46) = 12.75, *p* < .001, 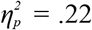. Simple effects analyses of the interaction indicated that differential responding (CS+ > CS−) remained significant in the DR group, *p* < .001, 95% CI [1.00, 1.92], whereas no significant difference between the CS+ and the CS− was observed in the Ctrl group, *p* = .175. In addition, US expectancy ratings for the CS+ were significantly higher in the DR group than in the Ctrl group, *p* < .001, 95% CI [−2.76, −0.90], whereas the two groups did not differ significantly in US expectancy for the CS−, *p* = .110.

For the arousal and valence ratings collected after re-extinction, the main effect of Stimulus was significant for arousal (CS+ > CS−), *F*(1, 46) = 21.36, *p* < .001, 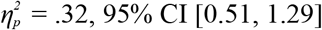, whereas the main effect of Group was not significant, *F*(1, 46) = 2.19, *p* = .146, and the Group × Stimulus interaction was not significant, *F*(1, 46) = 1.95, *p* = .169. For valence, the main effect of Stimulus was significant (CS+ > CS−), *F*(1, 46) = 9.71, *p* = .003, 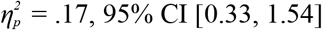, whereas the main effect of Group was not significant, *F*(1, 46) = 0.13, *p* = .724, and the Group × Stimulus interaction was not significant, *F*(1, 46) = 2.54, *p* = .118.

### 3.4. Manipulation Checks

Accuracy on the restricted retrieval trials was analyzed in the DR group (n = 24) to verify compliance with the 1-back instruction. Because no US was delivered during either extinction phase, the correct judgment on every restricted retrieval trial was “safe.” Accuracy was high overall (M = 92.69%, SD = 12.54) and significantly above the chance level of 50%, *t*(23) = 16.67, *p* < .001, *d* = 3.40. Accuracy was also significantly above chance in each condition separately, all *ts* > 5.79, all *ps* < .001 (Bonferroni-corrected), and all 24 participants individually performed above chance (range 54.69%–100.00%). A 2 (Day: Day 2, Day 3) × 2 (Stimulus: CS+, CS−) repeated-measures ANOVA revealed a significant main effect of Stimulus, *F*(1, 23) = 4.78, *p* = .039, 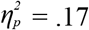, with lower accuracy for the CS+ (M = 87.54%, SD = 23.32) than for the CS− (M = 97.84%, SD = 6.12), whereas the main effect of Day was marginal, *F*(1, 23) = 4.18, *p* = .053, 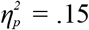(Day 2: M = 90.36%, SD = 15.58; Day 3: M = 95.02%, SD = 11.58), and the Day × Stimulus interaction was not significant, *F*(1, 23) = 0.69, *p* = .416. Because accuracy approached the ceiling in several conditions (13 of 24 participants responded correctly on every restricted retrieval trial), these effects were re-examined using Wilcoxon signed-rank tests, which confirmed the Stimulus effect (*p* = .023) but not the Day effect (*p* = .091). Adding Half (early vs. late trials) as a third within-subjects factor revealed neither a main effect of Half, *F*(1, 23) = 1.78, *p* = .195, 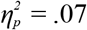, nor any interaction involving Half, all *Fs* < 1.86, all *ps* > .18, indicating that compliance did not change across trials within a phase.

Errors were highly informative with respect to the nature of participants’ responses. Across both extinction phases, 117 of 1,632 restricted retrieval trials (7.17%) were answered incorrectly, and 100 of these errors (85.5%) occurred on CS+ trials (CS+: 15.89% and 9.03% errors on Days 2 and 3; CS−: 3.39% and 0.93%). Thirteen participants made no errors at all, and five made errors only on CS+ trials. This asymmetry indicates that participants were not responding indiscriminately: the residual threat expectancy attached to the CS+ made it more difficult to register that this stimulus, too, had been safe on the preceding presentation.

Averaged across participants, mean response time on correct restricted retrieval trials was 2.69 s (SD = 0.84 s; median trial-level RT = 1.88 s), indicating that participants completed the judgment within a few seconds on each trial.

Taken together, these results indicate that participants in the DR group understood and complied with the 1-back instruction and consistently registered the absence of the US on the preceding trial.

### 3.5. Renewal Test

A Group × Stimulus × Trial (Trial 1, Trial 2) repeated-measures ANOVA revealed a significant main effect of Group (Ctrl > DR), *F*(1, 46) = 10.57, *p* = .002,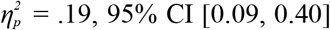; a significant main effect of Stimulus (CS+ > CS−), *F*(1, 46) = 13.29, *p* < .001, 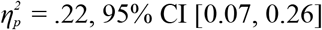; and a significant main effect of Trial (Trial 1 > Trial 2), *F*(1, 46) = 10.81, *p* = .002, 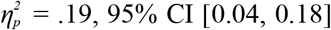. The Group × Stimulus interaction, *p* = .407; the Group × Trial interaction, *p* = .862; the Stimulus × Trial interaction, *p* = .815; and the Group × Stimulus × Trial interaction, *p* = .390, were all non-significant.

The same ANOVA was applied to the US expectancy, arousal, and valence ratings. For US expectancy, the main effect of Stimulus was significant (CS+ > CS−), *F*(1, 46) = 52.11, *p* < .001, 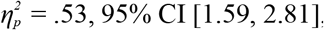, and the main effect of Trial was significant (Trial 1 > Trial 2), *F*(1, 46) = 11.62, *p* = .001, 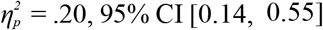, whereas the main effect of Group was not significant, *F*(1, 46) = 1.74, *p* = .193. Only the Group × Stimulus interaction was significant, *F*(1, 46) = 4.65, *p* = .036,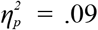. Simple effects analyses indicated that US expectancy ratings for the CS+ were significantly higher in the Ctrl group than in the DR group, *p* = .023, whereas the two groups did not differ significantly in US expectancy for the CS−, *p* = .766. The Group × Trial interaction, *p* = .608; the Stimulus × Trial interaction, *p* = .458; and the Group × Stimulus × Trial interaction, *p* = .750, were all non-significant.

For arousal, the main effect of Group was significant (Ctrl > DR), *F*(1, 46) = 5.81, *p* = .020, 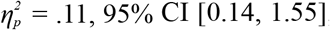, and the main effect of Stimulus was significant (CS+ > CS−), *F*(1, 46) = 46.71, *p* < .001, 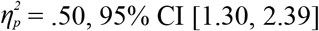, whereas the main effect of Trial was not significant, *F*(1, 46) = 1.85, *p* = .181. The Group × Stimulus interaction, *p* = .183; the Group × Trial interaction, *p* = .352; the Stimulus × Trial interaction, *p* = .072; and the Group × Stimulus × Trial interaction, *p* = .072, were all non-significant.

For valence, the main effect of Group was significant (Ctrl > DR), *F*(1, 46) = 4.84, *p* = .033, 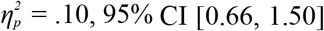, and the main effect of Stimulus was significant (CS+ > CS−), *F*(1, 46) = 20.88, *p* < .001, 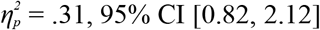, whereas the main effect of Trial was not significant, *F*(1, 46) = 1.48, *p* = .229. The Group × Stimulus interaction, *p* = .923; the Group × Trial interaction, *p* = .348; the Stimulus × Trial interaction, *p* = .122; and the Group × Stimulus × Trial interaction, *p* = .204, were all non-significant.

Both the SCR and subjective rating results indicate that fear renewal was weaker in the DR group than in the Ctrl group.

## 4. Discussion

Drawing on the distinct modulatory effects of selective retrieval on competing memories and on the learning-enhancing effects of variability, we developed a novel dual-retrieval-mode extinction training procedure that combines unrestricted retrieval with restricted 1-back retrieval, and compared it with standard extinction, which relies on a single unrestricted retrieval mode. Conditioned responding was indexed throughout using skin conductance responses together with US-expectancy, valence, and arousal ratings. The results showed that, relative to standard extinction training, dual-retrieval-mode extinction training reduced fear renewal. This finding provides preliminary evidence that varying memory retrieval demands during extinction may enhance the contextual generalizability of extinction learning. Importantly, this procedure was not designed to target a single theoretical mechanism; rather, it was intended to integrate several potentially beneficial features identified in prior learning and memory research, including selective retrieval, awareness of the CS safety outcome, and variability in retrieval mode.

### 4.1. Reduced Habituation and Fear Renewal Under the Dual-Retrieval Mode

At the end of fear acquisition, both SCRs and all subjective measures confirmed comparable levels of conditioned fear in the two groups, indicating that the groups were well matched prior to the experimental manipulation. During extinction and re-extinction, however, the different retrieval modes gave rise to marked between-group differences in SCRs and US expectancy ratings in the later part of extinction: the DR group showed significantly higher SCRs and US expectancy ratings than the Ctrl group, and significant differential US expectancy (CS+ > CS−) was maintained only in the DR group, whereas the Ctrl group showed no difference between the CS+ and the CS− in US expectancy by the end of each extinction phase. At first glance, these findings appear inconsistent with the reduction in fear renewal observed in the DR group on Day 4. However, the post-extinction arousal and valence ratings revealed no differences between the Ctrl group and the DR group, and the Ctrl group continued to show a significant difference between the CS+ and the CS−. This dissociation suggests that the reduced SCRs and US expectancy ratings to the CS+ on the final two trials of extinction in the Ctrl group may reflect within-session habituation rather than stronger inhibitory learning, because the process of extinction involves both inhibitory learning and habituation (Brown et al., 2017; Craske et al., 2014). The Ctrl group performed the same US expectancy rating on every trial throughout extinction, and this repetitive, uniform, and highly predictable response pattern would be expected to promote habituation. In the DR group, by contrast, the final two trials of each extinction phase required switching from the safe/dangerous judgment used under the restricted 1-back retrieval mode to the US expectancy rating used under the unrestricted retrieval mode, and this transition eliminated the habituated response. This interpretation is further supported by the results of the renewal test on Day 4, in which all outcome measures (SCR, US expectancy, arousal, and valence) consistently indicated weaker conditioned fear to the CS+ in the DR group than in the Ctrl group. These findings are consistent with previous research showing that performance at the end of extinction is a poor predictor of long-term treatment outcome (Cooper et al., 2017), and they highlight the importance of distinguishing performance during extinction from the durability of extinction memory.

One finding that departs from our predictions warrants comment. At the end of re-extinction, SCRs in the DR group were higher to the CS− than to the CS+ (p = .009), whereas the Ctrl group showed no difference between the two stimuli (p = .791). This pattern should be interpreted cautiously. It emerged against a background of generally elevated SCRs in the DR group, which we attribute to reduced within-session habituation, and the absolute level of responding at the end of an extinction phase is a poor predictor of long-term outcome (Cooper et al., 2017). Critically, the pattern did not persist: at the renewal test, the DR group again showed robust differential responding, with higher SCRs, US expectancy, arousal, and negative valence for the CS+ than for the CS−, indicating that the original CS–US association remained intact. The end-of-re-extinction reversal is therefore likely to reflect transient differences in habituation and in task-related arousal rather than a reversal of the underlying associative structure.

It is also worth noting that the reduction was CS+-specific for US expectancy, whereas for SCRs, arousal, and valence the Group main effect was not qualified by an interaction, indicating a broader reduction in reactivity to both stimuli. This pattern suggests that the dual-retrieval mode enhanced the accessibility of the extinction memory rather than selectively down-regulating the CS+ alone.

### 4.2. Possible Mechanisms Underlying the Dual-Retrieval-Mode Effect

The attenuation of fear renewal may stem from several beneficial features of the dual-retrieval mode. The first concerns selective retrieval itself. Restricted retrieval in the 1-back format requires participants to recall the safety outcome of the preceding trial selectively before judging the outcome of the current CS, and then to make a judgment consistent with that recollection. Previous RP and RIF research has shown that selectively retrieving one memory representation can simultaneously strengthen that memory (the RP effect; Karpicke & Blunt, 2011) and reduce the accessibility of competing representations that share the same retrieval cue (the RIF effect; Anderson et al., 1994; Murayama et al., 2014). Accordingly, repeatedly retrieving the safety-related CS– no US association in the present study may both increase the strength and accessibility of that memory (RP effect) and suppress retrieval of the CS–US association, thereby removing the proactive interference of the fear memory with extinction learning. Because the extinction memory and the fear memory are mutually exclusive competitors, strengthening the extinction memory may in turn produce retroactive interference with the fear memory (RIF effect).

It should be noted, however, that this account does not imply that the fear memory was erased. During the renewal test, the DR group still showed significantly stronger responses to the CS+ than to the CS− in SCR, US expectancy, arousal, and valence, indicating that the original CS–US association remained intact. The attenuation of fear renewal is therefore unlikely to reflect forgetting or erasure of the fear memory itself. Rather, these findings are more consistent with the interpretation that the dual-retrieval mode increased the relative accessibility or competitive strength of the extinction memory, enabling it to compete more effectively with the fear memory and also to suppress the expression of the fear memory. From both evolutionary and clinical perspectives, such an outcome may be preferable to the complete erasure of the fear memory: the ability to improve the regulation of fear expression while preserving an adaptive representation of threat may be more ecologically functional and clinically realistic than eliminating the fear memory altogether (Craske, Sandman, and Stein, 2022).

A second feature concerns awareness of the shift in contingency. Restricted 1-back retrieval maintained participants’ attention to trial-by-trial changes in contingency and promoted awareness of the shift in contingency from CS–US to CS–no US, facilitating memory updating (Jovanovic et al., 2022). Direct behavioral support for this claim comes from the manipulation checks: overall accuracy on the restricted retrieval trials was high (92.69%), but accuracy was significantly lower for the CS+ than for the CS−, and 85.5% of all errors occurred on CS+ trials. If participants had simply learned that “nothing ever happens” and responded “safe” indiscriminately, errors would have been distributed evenly across the two stimuli. The concentration of errors on the CS+ instead indicates that participants were processing each stimulus individually and that the residual threat expectancy attached to the CS+ made it specifically harder to register that this stimulus, too, had been safe on the preceding presentation.

A third feature concerns prediction error. The US expectancy rating trials conducted under the unrestricted retrieval mode preserved opportunities for prediction error. Although the DR group completed fewer unrestricted retrieval trials, for the updating of associative memory the cumulative magnitude of prediction error across an entire phase may matter more than the absolute number of trials involving prediction error (Chan & Harris, 2019). The Ctrl group, repeatedly exposed to the non-reinforced CS with an unchanging response mode, would show greatly reduced or absent prediction error after the first few trials, such that learning on later trials promoted habituation more than continued updating of associative memory. In the DR group, by contrast, the switch between retrieval modes itself elicited prediction errors.

Finally, the same switching between retrieval modes also increased variability. Variability in learning conditions is thought to enhance inhibitory learning by preventing rigid, repetition-based responding and by promoting more flexible and context-independent retrieval of the extinction memory (Craske et al., 2014). Alternating between retrieval modes may have broadened the range of retrieval cues associated with the CS–no US memory, making that memory more accessible across contexts. Because the dual-retrieval-mode extinction procedure manipulated these features simultaneously, their independent contributions cannot be isolated; these components may therefore act in a synergistic manner.

### 4.3. Differences From the Standard Retrieval Practice/RIF Paradigm

The present findings also highlight important differences between selective retrieval practice in conditioned fear and the typical RP/RIF paradigm. In typical retrieval practice/retrieval-induced forgetting studies, competing items associated with the same retrieval cue are generally comparable in memory strength, and the competing representations, although different, are not mutually exclusive (Reeck & LaBar, 2023; Ye et al., 2020). In fear conditioning, by contrast, a well-consolidated CS–US association is established before extinction begins, such that the competing CS–no US association is introduced against the background of a considerably stronger original fear memory. Moreover, the two associations are directly opposed to one another, a situation that has no counterpart in standard word-list RP/RIF paradigms. Testing in the typical RP/RIF paradigm is also qualitative in nature: given the cue Fruit– at test, participants either recall Orange and Pear or fail to recall them. In fear-conditioning research, in order to capture the ongoing and dynamic competition between two opposing memory traces, trial-by-trial US expectancy ratings are typically used instead of categorical recall measures. This quantitative approach permits a fine-grained assessment of changes in the relative strength of competing memories throughout the learning process.

### 4.4. Limitations and Future Directions

This study has several limitations. First, although the dual-retrieval-mode extinction procedure was theoretically informed by retrieval practice, retrieval-induced forgetting, prediction error, and variability-based learning, the present design cannot isolate the contribution of any individual component. One next step is to include additional control conditions to determine the unique contribution of selective retrieval independently of prediction error and variability. One such condition would be a pure restricted retrieval condition without alternation between retrieval modes. Second, we used a laboratory fear-conditioning paradigm with healthy participants; whether the dual-retrieval-mode extinction procedure generalizes to threat stimuli with greater ecological validity, or to clinical populations with anxiety and trauma-related disorders, remains to be examined. Third, beyond fear renewal, other forms of fear relapse exist, such as spontaneous recovery and reinstatement; future research should examine the effects of the dual-retrieval-mode extinction procedure on these other forms of fear relapse. Fourth, the sample consisted predominantly of female participants (41 female, 7 male) drawn from a single university population and a narrow age range (18–21 years), all of whom were of the same ethnic background. The sex imbalance reflects the gender composition of the psychology student pool from which participants were recruited.

## 5. Conclusion

This study found that an extinction training procedure combining unrestricted retrieval and selective retrieval within a dual-retrieval mode reduced fear renewal relative to a standard extinction procedure. This suggests that strategically manipulating retrieval demands during extinction may be a promising and theoretically grounded avenue for enhancing the durability and contextual generalizability of extinction learning. More broadly, these findings highlight retrieval mode as a potentially important yet underexplored dimension of extinction learning that may complement existing approaches to optimizing exposure therapy. Future research should determine the specific contributions of the individual procedural components and evaluate the efficacy of this approach in clinical populations and for other forms of fear relapse.

## CRediT authorship contribution statement

Yingzhu Zeng: Conceptualization, Data curation, Formal analysis, Writing – original draft, Writing – review & editing.

Zimeng Li: Methodology, Investigation, Data curation.

Hongbo Wang: Conceptualization, Methodology, Formal analysis, Writing – original draft, Writing – review & editing, Supervision, Project administration, Funding acquisition.

## Data Availability

The data that support the findings of this study are available from the corresponding author upon reasonable request.

## Declaration of generative AI and AI-assisted technologies in the manuscript preparation process

During the preparation of this work the authors used Yuanbao (Tencent Large Model, Shenzhen, China; https://yuanbao.tencent.com/) for English translation and language editing. After using this tool, the authors reviewed and edited the content as needed and take full responsibility for the content of the published article.

## References

Anderson, M. C. (2003). Rethinking interference theory: Executive control and the mechanisms of forgetting. Journal of Memory and Language, 49(4), 415–445. 10.1016/j.jml.2003.08.006

Anderson, M. C., Bjork, R. A., & Bjork, E. L. (1994). Remembering can cause forgetting: retrieval dynamics in long-term memory. Journal of Experimental Psychology: Learning, Memory, and Cognition, 20(5), 1063–1087. 10.1037/0278-7393.20.5.1063

Anderson, M. C., & Hulbert, J. C. (2021). Active Forgetting: Adaptation of Memory by Prefrontal Control. Annual Review of Psychology, 72, 1–36. 10.1146/annurev-psych-072720-094140

Bjork, R. A., & Bjork, E. L. (2006). Optimizing treatment and instruction: Implications of a new theory of disuse. In L.-G. Nilsson & N. Ohta (Eds.), Memory and society: Psychological perspectives (pp. 109–133). Psychology Press.

Bouton, M. E. (1993). Context, time, and memory retrieval in the interference paradigms of Pavlovian learning. Psychological Bulletin, 114(1), 80–99. 10.1037/0033-2909.114.1.80

Bouton, M. E., & Moody, E. W. (2004). Memory processes in classical conditioning. Neuroscience & Biobehavioral Reviews, 28(7), 663–674. 10.1016/j.neubiorev.2004.09.001

Brown, L. A., LeBeau, R. T., Chat, K. Y., & Craske, M. G. (2017). Associative learning versus fear habituation as predictors of long-term extinction retention. Cognition and Emotion, 31(4), 687–698. 10.1080/02699931.2016.1158695

Carpenter, J. K., Andrews, L. A., Witcraft, S. M., Powers, M. B., Smits, J. A. J., & Hofmann, S. G. (2018). Cognitive behavioral therapy for anxiety and related disorders: A meta-analysis of randomized placebo-controlled trials. Depression and Anxiety, 35(6), 502–514. 10.1002/da.22728

Chan, C. K. J., & Harris, J. A. (2019). The partial reinforcement extinction effect: The proportion of trials reinforced during conditioning predicts the number of trials to extinction. Journal of Experimental Psychology: Animal Learning and Cognition, 45(1), 43–58. 10.1037/xan0000190

Chen, T., Mei, Y., Zhou, S., Dou, H., & Lei, Y. (2024). Trait self-compassion enhances activation in the medial prefrontal cortex during fear extinction: An fNIRS study. International Journal of Clinical and Health Psychology, 24(4), 100516. 10.1016/j.ijchp.2024.100516

Chen, W., Li, J., Zhang, X., Dong, Y., Shi, P., Luo, P., & Zheng, X. (2021). Retrieval-extinction as a reconsolidation-based treatment for emotional disorders: Evidence from an extinction retention test shortly after intervention. Behaviour Research and Therapy, 139, 103831. 10.1016/j.brat.2021.103831

Cicchini, G. M., Mikellidou, K., & Burr, D. C. (2024). Serial Dependence in Perception. Annual Review of Psychology, 75, 129–154. 10.1146/annurev-psych-021523-104939

Cooper, A. A., Clifton, E. G., & Feeny, N. C. (2017). An empirical review of potential mediators and mechanisms of prolonged exposure therapy. Clinical Psychology Review, 56, 106–121. 10.1016/j.cpr.2017.07.003

Craske, M. G., Sandman, C. F., & Stein, M. B. (2022). How can neurobiology of fear extinction inform treatment? Neuroscience & Biobehavioral Reviews, 143, 104923. 10.1016/j.neubiorev.2022.104923

Craske, M. G., Treanor, M., Conway, C. C., Zbozinek, T., & Vervliet, B. (2014). Maximizing exposure therapy: an inhibitory learning approach. Behaviour Research and Therapy, 58, 10–23. 10.1016/j.brat.2014.04.006

Craske, M. G., Treanor, M., Zbozinek, T. D., & Vervliet, B. (2022). Optimizing exposure therapy with an inhibitory retrieval approach and the OptEx Nexus. Behaviour Research and Therapy, 152, 104069. 10.1016/j.brat.2022.104069

Dudziak, M., Franssen, M., Vervliet, B., & Beckers, T. (2025). The impact of losing control over threat on the acquisition, extinction, and renewal of conditioned fear. Behaviour Research and Therapy, 193, 104842. 10.1016/j.brat.2025.104842

Faul, F., Erdfelder, E., Buchner, A., & Lang, A. G. (2009). Statistical power analyses using G*Power 3.1: tests for correlation and regression analyses. Behavior Research Methods, 41(4), 1149–1160. 10.3758/brm.41.4.1149

Glazier, B. L., Alden, L. E., & Graf, P. (2021). Retrieval-induced forgetting in a social task. Cognition and Emotion, 35(1), 199–206. 10.1080/02699931.2020.1806042

JASP Team. (2025). JASP (Version 0.95.4) [Computer software]. https://jasp-stats.org/

Jovanovic, T., Wiltshire, C. N., Reda, M. H., France, J., Wanna, C. P., Minton, S. T., Davie, W., Grasser, L. R., Winters, S., Schacter, H., Marusak, H. A., & Stenson, A. F. (2022). Uncertain in the face of change: Lack of contingency shift awareness during extinction is associated with higher fear-potentiated startle and PTSD symptoms in children. International Journal of Psychophysiology, 178, 90–98. 10.1016/j.ijpsycho.2022.06.008

Karpicke, J. D., & Blunt, J. R. (2011). Retrieval practice produces more learning than elaborative studying with concept mapping. Science, 331(6018), 772–775. 10.1126/science.1199327

Kim, S., Burr, D., Cicchini, G. M., & Alais, D. (2020). Serial dependence in perception requires conscious awareness. Current Biology, 30(6), R257–R258. 10.1016/j.cub.2020.02.008

Lonsdorf, T. B., Menz, M. M., Andreatta, M., Fullana, M. A., Golkar, A., Haaker, J., Heitland, I., Hermann, A., Kuhn, M., Kruse, O., Meir Drexler, S., Meulders, A., Nees, F., Pittig, A., Richter, J., Römer, S., Shiban, Y., Schmitz, A., Straube, B.,… Merz, C. J. (2017). Don’t fear ‘fear conditioning’: Methodological considerations for the design and analysis of studies on human fear acquisition, extinction, and return of fear. Neuroscience & Biobehavioral Reviews, 77, 247–285. 10.1016/j.neubiorev.2017.02.026

Loock, K., Kalbe, F., & Schwabe, L. (2025). Cognitive mechanisms of aversive prediction error-induced memory enhancements. Journal of Experimental Psychology: General, 154(4), 1102–1121. 10.1037/xge0001712

Murayama, K., Miyatsu, T., Buchli, D., & Storm, B. C. (2014). Forgetting as a consequence of retrieval: a meta-analytic review of retrieval-induced forgetting. Psychological Bulletin, 140(5), 1383–1409. 10.1037/a0037505

Myers, K. M., & Davis, M. (2007). Mechanisms of fear extinction. Molecular Psychiatry, 12(2), 120–150. 10.1038/sj.mp.4001939

O’Malley, K. R., & Waters, A. M. (2018). Attention avoidance of the threat conditioned stimulus during extinction increases physiological arousal generalisation and retention. Behaviour Research and Therapy, 104, 51–61. 10.1016/j.brat.2018.03.001

Ortega, J., Chen, Z., & Whitney, D. (2023). Serial dependence in emotion perception mirrors the autocorrelations in natural emotion statistics. Journal of Vision, 23(3), 12. 10.1167/jov.23.3.12

Pace-Schott, E. F., Tracy, L. E., Rubin, Z., Mollica, A. G., Ellenbogen, J. M., Bianchi, M. T., Milad, M. R., Pitman, R. K., & Orr, S. P. (2014). Interactions of time of day and sleep with between-session habituation and extinction memory in young adult males. Experimental Brain Research, 232(5), 1443–1458. 10.1007/s00221-014-3829-9

Reeck, C., & LaBar, K. S. (2023). Retrieval-induced forgetting of emotional memories. Cognition and Emotion, 1–17. 10.1080/02699931.2023.2279156

Roediger, H. L., 3rd, & Butler, A. C. (2011). The critical role of retrieval practice in long-term retention. Trends in Cognitive Sciences, 15(1), 20–27. 10.1016/j.tics.2010.09.003

Smith, A. M., Floerke, V. A., & Thomas, A. K. (2016). Retrieval practice protects memory against acute stress. Science, 354(6315), 1046–1048. 10.1126/science.aah5067

Sperl, M. F. J., Wroblewski, A., Mueller, M., Straube, B., & Mueller, E. M. (2021). Learning dynamics of electrophysiological brain signals during human fear conditioning. NeuroImage, 226, 117569. 10.1016/j.neuroimage.2020.117569

Stemerding, L. E., van Ast, V. A., & Kindt, M. (2023). Manipulating expectancy violations to strengthen the efficacy of human fear extinction. Behaviour Research and Therapy, 165, 104319. 10.1016/j.brat.2023.104319

Stussi, Y., Ferrero, A., Pourtois, G., & Sander, D. (2019). Achievement motivation modulates Pavlovian aversive conditioning to goal-relevant stimuli. npj Science of Learning, 4(1), 4. 10.1038/s41539-019-0043-3

Vervliet, B., Craske, M. G., & Hermans, D. (2013). Fear extinction and relapse: state of the art. Annual Review of Clinical Psychology, 9, 215–248. 10.1146/annurev-clinpsy-050212-185542

Wang, Y., Zhu, Z., Hu, J., Schiller, D., & Li, J. (2021). Active suppression prevents the return of threat memory in humans. Communications Biology, 4(1), 609. 10.1038/s42003-021-02120-2

Wicking, M., Steiger, F., Nees, F., Diener, S. J., Grimm, O., Ruttorf, M., Schad, L. R., Winkelmann, T., Wirtz, G., & Flor, H. (2016). Deficient fear extinction memory in posttraumatic stress disorder. Neurobiology of Learning and Memory, 136, 116–126. 10.1016/j.nlm.2016.09.016

Ye, Z., Shi, L., Li, A., Chen, C., & Xue, G. (2020). Retrieval practice facilitates memory updating by enhancing and differentiating medial prefrontal cortex representations. eLife, 9. 10.7554/eLife.57023

